# Extracellular Vesicles Mediate Activation and Trafficking of Splenic Immune Cells to the Heart Post-Myocardial Infarction

**DOI:** 10.64898/2026.06.23.734125

**Authors:** Kaneez Fatima, Austin Angelotti, Vinay Kumar, Vishnu Chollangi, Wafa Aziz, Siddarth Dasari, Emily N. Bianchini, Joseph Wang, Suman Asalla, Harpreet Singh, Sumanth D. Prabhu, Shyam S. Bansal

## Abstract

**Background:** Myocardial infarction (MI) triggers splenic immune cell trafficking to the heart. Vehicles that carry these signals and mediate this crosstalk are unknown.

**Hypothesis:** We hypothesize that extracellular vesicles (EVs) released post-MI mediate splenic immune trafficking to the heart.

**Methods:** Mice were treated daily with an EV biogenesis inhibitor (GW4869) or vehicle. Splenic/cardiac immune cells were assessed at 3d while survival, cardiac function, hypertrophy, and fibrosis were evaluated at 8w post-MI. Plasma EVs from 1d MI mice or from the hearts that underwent MI/sham in a Langendorff system induced splenic immune trafficking to the heart within 3d and systolic dysfunction at 8w in naïve mice.

**Results:** GW4869 i) inhibited splenic regression, ii) increased splenic retention of neutrophils, monocytes, dendritic cells (DCs), and CD4⁺ T-cells, iii) decreased cardiac gene expression of pro-inflammatory cytokines/chemokines, and iv) decreased trafficking of immune cells to the hearts at 3d post-MI, and iii) improved systolic function and attenuated hypertrophy at 8w post-MI. MI EVs accumulated in the spleen and promoted egress of matured splenic immune cells upon administration to naïve mice. Cardiac pro-inflammatory cytokines/chemokines expression and CCR2^+^MHC-II^hi^ infiltrating macrophages, CD11c^+^ DCs, and CD4^+^ and CD4^+^TNFα^+^ T-cell levels were also increased in naïve mice at 3d post-injection. Importantly, transfer of MI EVs for 2 days induced systolic dysfunction, cellular hypertrophy, and fibrosis in naïve mice at 8 w post-injection. DCs process MI EVs for T-cells activation.

**Conclusions:** EVs mobilize splenic immune cells to the heart post-MI and their inhibition can subdue inflammatory tissue-damage to promote healing post-MI.

## Introduction

During acute myocardial infarction (MI) dying cardiomyocytes (CMs) and degradation of the extracellular matrix in the infarcted tissue release a broad range of alarmins, called damage-associated molecular patterns (DAMPs), such a**s** HMGB1, ATP, heat shock proteins, and calreticulin^1^. Circulating DAMPs act as endogenous danger signals to activate pattern recognition receptors (PRRs), and set in motion a complex signaling cascade ^2^ designed to recruit innate and adaptive immune cells^2–6^ from the hematopoietic and extra-hematopoietic tissues such as the bone marrow (BM)^6,7^ and the spleen^8,9^ for tissue-repair. Despite extensive evidence supporting the role of DAMPs in immune activation, the mechanisms by which these signals are transported from the injured myocardium to the spleen to activate and recruit immune cells are unknown and is a big constraint in our understanding of immune activation mechanisms post-ischemic cardiac injury.

Recently, extracellular vesicles (EVs) have been demonstrated to play an important role in facilitating antigen presentation and regulating immune activation in the spleen and other tissues^10^. Almost all cells, including CMs, endothelial cells, vascular smooth muscle cells, fibroblasts, and immune cells, release 40-120 nm EVs to shuttle bioactive molecules such as microRNAs (miRNAs), messenger RNAs (mRNAs), long non-coding RNAs (lncRNAs), proteins, DNA fragments, and lipids to effect intercellular juxtacrine and paracrine communication to convey signals for physiological homeostasis and/or stress^11–13^. These nanoscale vesicles are internalized by the recipient cells to modulate gene expression and cellular behavior through the processing of bioactive cargo contained within. Proteomic profiling of EVs show a diverse repertoire of immunomodulatory molecules, including cytokines, chemokines, and their receptors (IL-10, CCL2, VEGF-C, C-reactive protein) which can modulate immune cell function during disease states^13,14^. Exosomal miRNAs such as miR-155 and miR-21 have also been shown to regulate key inflammatory pathways, including NF-κB signaling, thereby controlling cytokine production and immune cell activation^15–17^. Additionally, it has been shown that protein cargo in EVs released by the hypoxic CMs contain cell-specific stress signals such as HSP60, HSP90, HSP27, Myosin-binding protein C, αβ-Crystallin, Tropomyosin-α1, cardiac muscle Actin and functional ion channels, that can further regulate immune cell activation and trafficking during ischemic cardiac injury^18^.

Since EVs are formed in the cytoplasm and contain a part of the parent cell’s plasma membrane, it is possible that cytosolic or membranous proteins from apoptotic cardiomyocytes are carried by the EVs, in the form of DAMPs, to facilitate antigen-presentation and immune activation post-MI. Thus, we hypothesized that EVs released from the stressed and dying CMs act as DAMP carriers to mediate paracrine activation of the immune responses by facilitating communication between the injured heart and the spleen. In this study, we demonstrate a critical role of EVs in the activation of splenic immune cells and their trafficking to the ischemic heart. We also show that concentration and protein cargo of EVs released by the infarcted heart are significantly different than the EVs from the sham hearts. We further investigated preferential localization of MI EVs in the spleens and their potential to induce systolic dysfunction and LV remodeling via splenic immune activation. Our findings suggest that EVs released by the infarcted heart carry injury-signals required for the activation and trafficking of splenic immune cells to the heart.

## Methods

### Rodent Studies

C57BL/6 mice (male, 10-12 week-old) used in the study were either purchased from Jackson Laboratory, USA or were bred in our facility. All experimental procedures were approved by the Institutional Animal Care and Use Committee of the Pennsylvania State College of Medicine (IACUC# PROTO202302485) and conducted in accordance with National Institutes of Health Guide for the Care and Use of Laboratory Animals (Department of Health and Human Services Publication No. 85-23, revised 1996). Mice were maintained under 12 h light/12 h dark cycle, with constant room temperature and humidity. All mice had free access to food and water ad libitum and 177 mice were used for all the studies.

### Myocardial Infarction Mouse Model and Surgical Procedure

10-12 week-old male mice were anesthetized using Isoflurane and underwent permanent left anterior descending coronary artery ligation (n=119) to induce MI, as described previously ^19^. Cardiac function was measured at 2 weeks and 8 weeks using echocardiography. At 3 days and 8 weeks, mice were euthanized by cervical dislocation under anesthesia and hearts and spleens were harvested. Tissues were weighed and processed either for mononuclear cell isolation or histological analysis. Tibia length was used to normalize all gravimetric data. Plasma was collected to measure EV levels.

### EV Biogenesis Inhibitor Treatment

All C57BL/6 mice were randomly divided into two groups to receive either vehicle or EV biogenesis inhibitor, GW4869, (catalog# 6741; Tocris Bioscience). Stock solution (10 mg/mL) for GW4869 was prepared in DMSO and stored at -20 °C. At the time of dosing, 60 µL of drug was mixed in 60 µL Tween-20 and 480 µL saline (100 µg/100 µL), and a volume equivalent to a dose of 5 mg/kg GW4869 (or vehicle) was administered *via* intraperitoneal injection. Dosing was started one day prior to the MI and continued for two more days post-MI.

### Echocardiography

Cardiac function in recipient mice was measured at baseline (day 0) and at 2 weeks and 8 weeks after treatment with GW4869 or adoptive transfer of EVs using echocardiography and their respective controls. Parasternal long-axis B-mode imaging was performed under 1% to 2% isoflurane anesthesia using VevoF2 imaging system equipped with the UHF57x transducer and adjustable heated rail system. Systolic function was analyzed by Vevo LAB 5.9.0 software^20^.

### Isolated Heart Perfusion by Langendorff Method and EV Isolation

Mice were administered heparin (*i.p.,*100 USP units) and anesthetized with 4% isoflurane. Hearts were excised and cannulated through the ascending aorta for *ex-vivo* perfusion using Langendorff system and perfused with warm (37°C) Krebs-Henseleit buffer to maintain their physiological function. Once stabilized (3-5 min), MI was induced by ligating permanent left anterior descending coronary artery. For sham surgery controls, the needle was passed through the cardiac wall without ligation. Approximately 100 mL retrograde oxygenated perfusate was collected at a controlled flow rate of ∼ 2mL/min from MI/sham hearts. Following perfusion, hearts were collected and sectioned into four sections and incubated in 1% triphenyltetrazolium chloride (TTC; catalog#, T8877; Sigma) for 20 min at 37 °C followed by fixing in 4% paraformaldehyde. After 24 h, heart sections were imaged and % infarct area was measured using ImageJ software. The collected perfusate (∼100 mL) was used for EV isolation using a combination of centrifugation and size exclusion chromatography (Qiagen exoEasy Maxi Kit, catalog#, 76064). Isolated EVs were characterized for their size, concentration and expression of EV specific markers, CD63-AF488 (FAB5417G, RD systems), CD81-AF488 (FAB4865G, RD systems), and CD9-AF488 (FAB5218G, RD systems), using NanoFCM system (NanoFCM Co. Ltd, UK).

### Adoptive Transfer of EVs

MI/sham EVs (180x10^6^) were intravenously injected twice/day (8 h apart) for 2 days into naïve mice to determine their functional activity. Cardiac function of recipient mice was measured at baseline, 2 weeks and 8 weeks using echocardiography. At 3 days and 8 weeks post-transfer, hearts and spleens were harvested from the recipient mice to measure innate and adaptive immune cells, and LV remodeling (fibrosis, hypertrophy and gene expression), respectively.

### Immune cell isolation, fixation and flow cytometric analysis

Immune cells from heart and spleen were isolated and fixed according to our previously published protocol^21^. The antibodies used in the study are listed in the supplemental Table 1.

### LV Fibrosis and Hypertrophy

Formalin-fixed LV tissues were embedded in paraffin and sectioned (5 µm thickness), deparaffinized, and rehydrated before staining. Masson’s trichome staining was performed to measure fibrosis in the border and remote zones. Alexa Fluor 488–conjugated wheat germ agglutinin (catalog# W11261; Invitrogen) and DAPI were used to label cardiomyocyte membranes and nuclei, respectively. The area of centrally nucleated cells was measured by ImageJ software.

### Tissue homogenization, RNA extraction, cDNA preparation, Real-Time Quantitative PCR

LV tissue (∼50 mg) was homogenized in 1 mL Trizol reagent (catalog# 15596018; Invitrogen) and RNA was extracted. RNA concentration was determined by NanoDrop (Accuris) and cDNA was synthesized using iScript cDNA synthesis kit (catalog# 1708891; BioRad). Quantitative real-time PCR was performed using iTaq Universal syber green supermix (catalog# 1725121; BioRad) with pre-designed and validated primers. β-actin or GAPDH was used as endogenous control and fold change in mRNA expression of targeted genes was measured. The primer sequences used in this study are listed in the supplemental Table 2.

### Statistical Analysis

Experimental data are shown as mean ± SD. Statistical analyses were performed using GraphPad Prism version 10.2.2. Groups were analyzed either by unpaired two-tailed student’s T test for 2-groups or ANOVA (one-way/two-way) with or without repeated measures for more than 2-groups where appropriate. Details of statistical test and sample sizes are provided in figure legends.

## Results

Acute ischemic injury, such as MI, induces trafficking of immune cells from the spleen^8,9,22^ and the BM^7^ to the heart and as a result of this immune cell reshuffling spleen loses significant weight as early as one day post MI^8,9^. Our data also shows approximately 30% loss in spleen weight (**Figure S1A**) which sustains for up to 3-days post MI (**Figure 1A**), peak time for the inflammatory immune response, presumably due to DAMPs being delivered to the distant immune rich tissues such as the spleen. However, the vehicles that mediate this inter-organ crosstalk during MI are not known.

**Figure 1.**
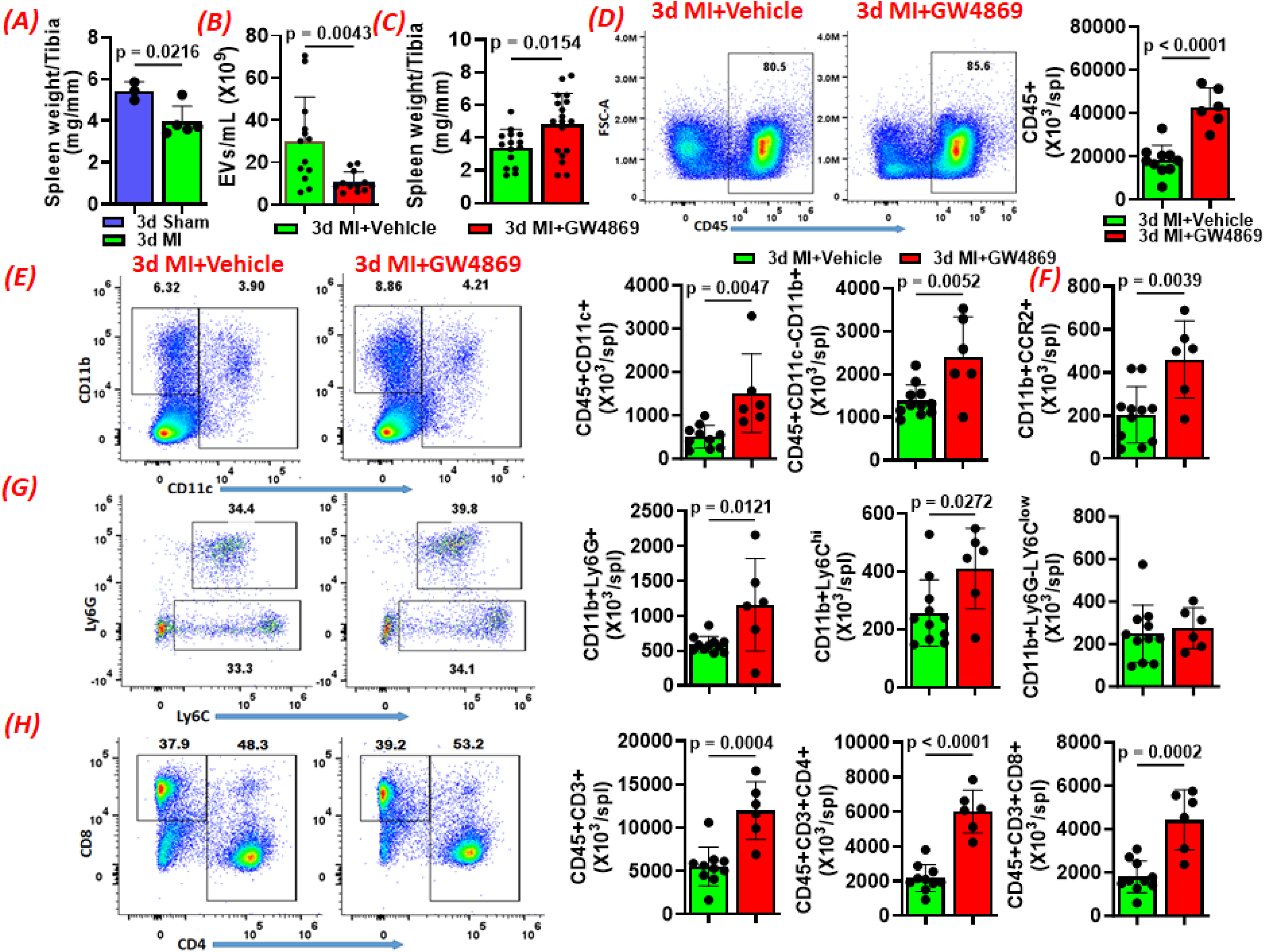
***(A)*** spleen weights in sham and MI mice at 3d post-surgery. ***(B)*** Plasma levels of extracellular vesicles (EVs) and ***(C)*** tibia normalized spleen weights in mice treated either with vehicle or EV biogenesis inhibitor, GW4869 (5 mg/kg; i.p.), at 3d post-MI. ***(D)*** Representative flow cytometry plots showing CD45^+^ leukocytes (left panel), and their group quantitation in the spleens of 3d MI mice treated either vehicle or GW4869. Representative flow cytometry plots and group quantitation for CD11c^+^ and CD11b^+^ myeloid cells ***(E)***, CD11b^+^CCR2^+^ cells ***(F)***, CD11b^+^Ly6C^+^ monocytes and CD11b^+^Ly6G^+^ leukocytes ***(G)***, and CD3^+^, CD3^+^CD4^+^ helper, and CD3^+^CD4^-^CD8^+^ cytotoxic T-cells in the spleens of 3d MI mice treated either with vehicle or GW4869. Each dot represents data from an individual mouse and all data were analyzed using two-tailed unpaired students T-test.

### EV Biogenesis Inhibition Blunts Splenic Regression and Immune Efflux Post-MI

To determine if EVs play a role in mediating splenic regression and immune efflux for their trafficking to the heart we administered (daily; i.p., 4 mg/kg) GW4869, an inhibitor of EV biogenesis, or vehicle to male C57BL/6 mice. Treatments were started a day before MI surgery and were continued daily until day 3 post-MI, as detailed in **Figuer S1B**. At day 3, mice were euthanized and their spleens and hearts were harvested and processed for immune cell analysis by flow cytometry using our previously described protocols^19,21^. GW4869 treatment significantly decreased EV release into the plasma at 3d post-MI (**Figure 1B**) without changing their size (**Figuer S1C**). Importantly, reduced EV biogenesis significantly blunted splenic regression at 3d post-MI as evident by increased tibia normalized spleen weight when compared with vehicle treated MI mice (**Figure 1C**). This was also reflected in the increased retention of splenic CD45^+^ leukocytes (**Figure 1D**). We further measured myeloid and lymphoid immune cells and found that EV biogenesis inhibitor significantly blunted the efflux resulting in the retention of both innate as well as adaptive immune cells from the spleen. Higher numbers of CD11c^+^ dendritic cells, CD11b^+^ myeloid cells, CD11b^+^CCR2^+^ cells, CD11b^+^Ly6G^+^ neutrophils and CD11b^+^Ly6C^hi^ monocytes (**Figure 1E, F, G**) as well as CD3^+^ T-cells, CD4^+^ helper T-cells (Th), and CD8^+^ cytotoxic T-cells (**Figure 1H**) were observed in the spleens of GW4869 treated mice, as compared to vehicle, at 3d post-MI.

### EV Biogenesis Inhibition Blunts Immune Trafficking to the Heart Post-MI

We also measured immune cells in the infarcted hearts at 3d post-MI after treatment with EV inhibitor as compared to vehicle. As shown in **Figure 2A**, treatment with GW4869 significantly reduced trafficking of CD45^+^ cells to the heart. A significant reduction in the innate immune cells such as CD11b^+^ myeloid cells, and among them CD11b^+^Ly6G^+^ neutrophils, CD11b^+^Ly6G^-^MHC-II^high^CCR2^+^ monocyte-derived macrophages (MDMs), and CD11b^+^Ly6G^-^MHC-II^low^CCR2^+^ infiltrating monocytes, including Ly6C^low^ (with a trending decrease in Ly6C^hi^) monocytes was also observed (**Figure 2B, C, and D)**. These changes in the innate immune cells were also complemented with reduced levels of CD4^+^ Th cells and its polarized subsets (**Figure 2E, F and G**). While pro-inflammatory IFNγ^+^ Th1 cells were significantly reduced, trending decreases in TNFα^+^, and IL-17^+^ Th17 cells were observed (**Figure 1E and F**). In contrast, anti-inflammatory subsets such as Th2 expressing IL-4 or regulatory T-cells expressing FoxP3 were not altered (**Figure 1F and 1G)** in GW4869 treated mice as compared to vehicle treated 3d MI mice. We also measured cardiac gene expression of different cytokines and chemokines known to mediate inflammatory immune responses post MI. As shown in **Figure 2H**, EV biogenesis inhibition also decreased gene expression of IL-1β, CCL-2, VEGF-c, and CTGF while IL-4, IL-6 and iNos showed decreasing trends, CD206, IL-10 TGF-β and collagen 1A1 and collagen 3A1 were not changed in hearts of mice treated with GW4869 when compared with vehicle treated 3d MI mice. These data suggest that EVs released by the infarcted hearts carry signals that are required for the activation and/or egress of the immune cells from the spleen and their trafficking to the ischemic heart.

**Figure 2.**
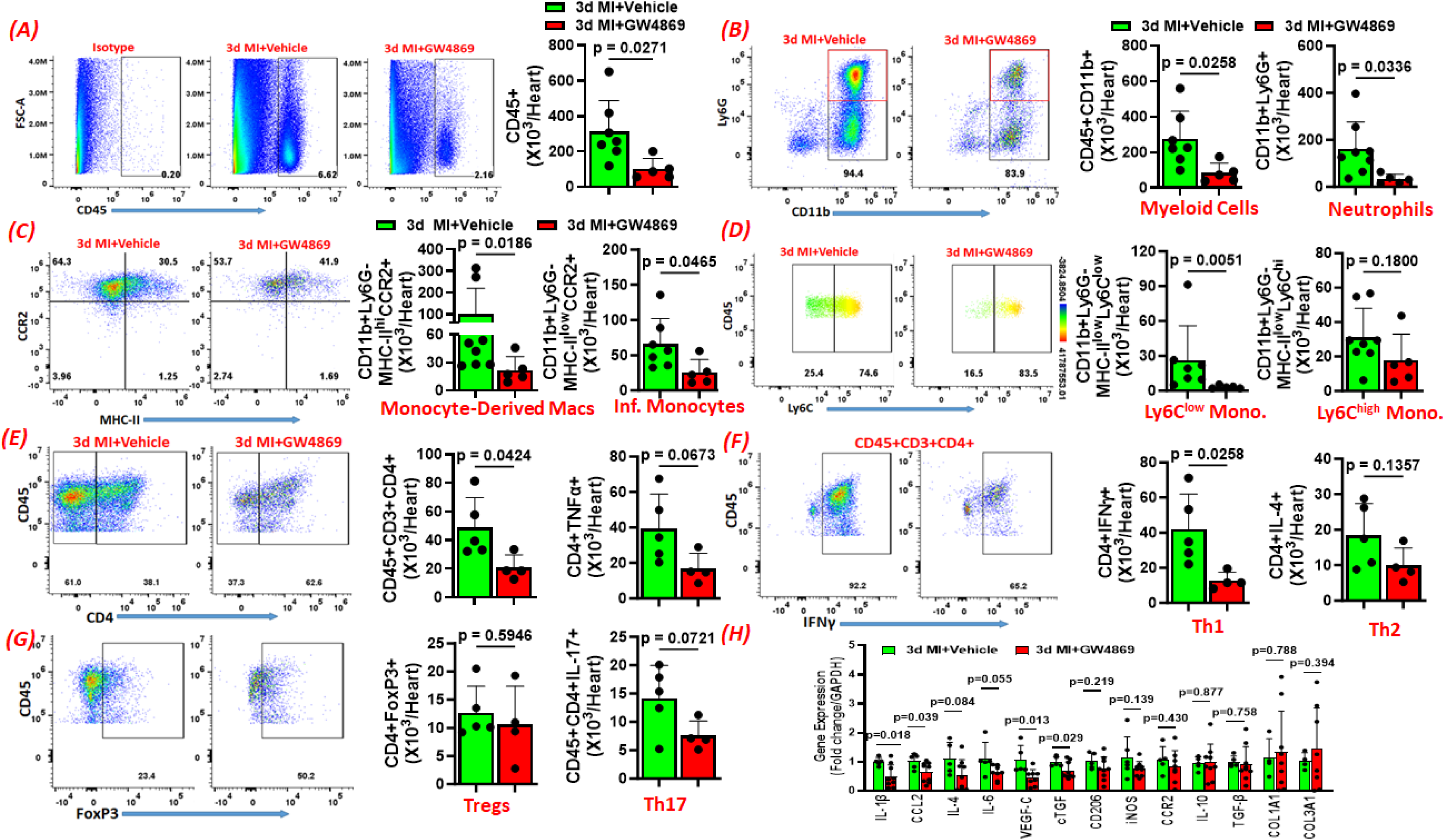
***(A)*** Representative flow cytometry plots showing CD45^+^ leukocytes (*left panel*), and their group quantitation (*right panel*) in the hearts of 3d MI mice treated either with vehicle or GW4869. ***(B)*** Representative flow cytometry plots and group quantitation for CD11b^+^ myeloid cells and CD11b^+^Ly6G^+^ neutrophils in the hearts of 3d MI mice treated either with vehicle or GW4869. ***(C)*** Representative flow cytometry plots and group quantitation for CD11b^+^MHC-II^hi^CCR2^+^ monocyte-derived macrophages and CD11b^+^MHC-II^low^CCR2^+^ monocytes in the hearts of 3d MI mice treated either with vehicle or GW4869. ***(D)*** Representative flow cytometry plots and group quantitation for CD11b^+^MHC-II^low^Ly6C^low^ anti-inflammatory and CD11b^+^MHC-II^low^Ly6C^hi^ pro-inflammatory monocytes in the hearts of 3d MI mice treated either with vehicle or GW4869. Representative flow cytometry plots and group quantitation for ***(E)*** CD45^+^CD3^+^CD4^+^ helper T-cells and CD4^+^TNFα^+^ helper T-cells, ***(F)*** CD4^+^IFNγ^+^ Th1 T-cells, and CD4^+^IL-4^+^ Th2 helper T-cells, and ***(G)*** CD4^+^FoxP3^+^ regulatory T-cells and CD4^+^IL-17^+^ Th17 T-cells in the hearts of 3d MI mice treated either with vehicle or GW4869. ***(H)*** Gene expression analysis of different cytokines and chemokines in the hearts of 3d MI mice treated either with vehicle or GW4869. Each dot represents data from an individual mouse and all data were analyzed using two-tailed unpaired students T-test.

### EV Biogenesis Inhibition Ameliorates Cardiac Hypertrophy and Dysfunction Post-MI

We further measured the effects of EV biogenesis inhibition during acute MI on overall cardiac function and LV remodeling during chronic HF (8 weeks post-MI). We treated MI mice with GW4869 (or vehicle) for 4 days starting a day before MI surgeries as before (**Figure S3A**) and measured their survival and changes in cardiac function over a period of 8 weeks. Inhibition of EV biogenesis, as compared to vehicle, did not result in increased survival (**Figure S2A**), but significantly improved cardiac function and decreased LV dilatation es evident by increased ejection fraction (EF) and decreased end-systolic and end-diastolic volumes (ESV and EDV) both at 2- and 8-weeks post-MI (**Figure 3B and C**). Heart rate was consistently maintained at >500 BPM in both the groups (**Figure S2B**). Although, tibia normalized heart weights (**Figure S2C**) or fibrosis **(Figure S2D)** was not different, significantly decreased cardiomyocyte hypertrophy was observed in GW4869 treated mice, as compared to vehicle treatment, consistent with decreased LV remodeling at 8 weeks post-MI **(Figure 3D)**. EV biogenesis inhibition and reduced immune trafficking also resulted in increased expression of angiogenic factors such as VEGF-R1 and VEGF-A (with a trending increase in IL-4) without altering pro-inflammatory cytokines or chemokines (**Figure S2E**).

**Figure 3.**
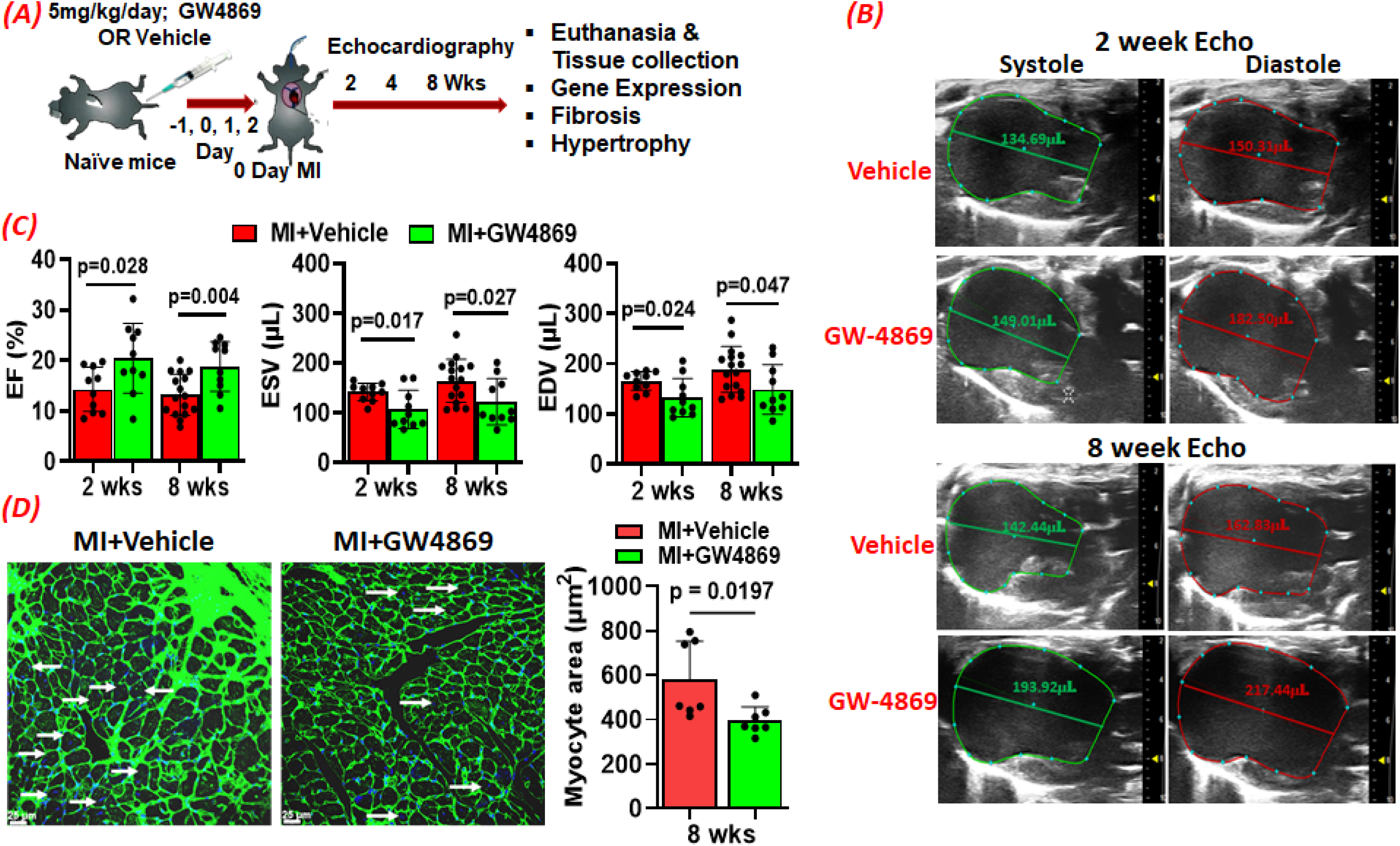
***(A)*** Schematic for the experimental plan to determine the effects of EV biogenesis inhibition in mediating cardiac dysfunction and LV remodeling post-MI in mice treated either with GW4869 or vehicle. ***(B)*** Representative parasternal B-mode images showing systole and diastole at 2 and 8 wks post-MI in mice treated either with the vehicle or GW4869. ***(C)*** group quantitation for the ejection fraction (EF), and end-systolic and end-diastolic volumes (ESV and EDV) at 2-and 8-wks post-MI in mice treated either with the vehicle or GW4869. ***(D)*** Representative photographs of LV sections stained with wheat germ agglutinin conjugated with fluorescein isothiocyanate (FITC; green) and DAPI (blue) to stain nuclei to show cardiomyocyte size (*left panel*) and group quantitation for cardiomyocyte area (*right panel*) at 8 wks post-MI in mice treated either with the vehicle or GW4869. Each dot represents data from a mouse and data in ***(C)*** and ***(D)*** were analyzed using two-tailed unpaired students T-test.

### Cardiac EVs Released in Response to Ischemic Cardiac Injury Induce Splenic Immune Activation and Trafficking to the Heart

To determine if EVs released from the infarcted hearts are sufficient to induce splenic immune activation, MI was induced in isolated hearts perfused using langendorff apparatus, 100 mL perfusate (∼50 mins) was collected, and EVs were isolated. Experimental protocol is shown in Fig 3A. Degree of infarction was confirmed using TTC staining and was found to be ∼40-50% (**Figure S3A and S3B**). Cryo-TEM analysis of isolated particles confirmed intact EV morphology with particles ranging from 60-120 nm **(Figure S3C**). EV release significantly increased immediately after MI without changes in their size (**Figure S3D**) or expression of EV markers such as CD9 or CD81 **(Figure S3E).** Importantly, protein cargo of MI EVs was significantly different than the sham EVs and was enriched in proteins such as complement factor D (CFD) and YWHAH known to attract and activate immune cells.

To determine if EVs generated post-MI localize to the spleen, 0.8X10^9^ EVs labeled with DiR dye were injected (i.v.; retro-orbital) into naïve mice and using IVIS their biodistribution was measured at different time intervals. While most of the sham EVs accumulated in the liver within one hour, MI EVs distributed to both the liver and the spleen (**Figure S4**). At 1d and 3d post-injection, we harvested different tissues and measured DiR intensity using IVIS. As shown in **Figure 4B**, most of the dye signal was localized in the liver and the spleen with little to no signal in other tissues including the brain, the heart, the lungs or the kidneys suggesting that EVs preferentially localize to the extrahematopoietic tissues. We further measured if injected EVs also activated splenic immune cells. To mimic MI like systemic milieu, we injected (retro-orbital) EVs two times a day 8 hours apart for 3 days (experimental design given in **Figure S5A**) and measured splenic immune activation at 3 days post-injection. As shown in **Figure 4C**, we observed significantly increased levels of antigen presenting myeloid cells (CD11b^+^ and CD11c^+^) and their expression of CCR2 and MHC-II, markers for increased trafficking and maturation, respectively (**Figure 4D and S5B**). We also observed increased levels of splenic CD11b^+^Ly6C^low^ monocytes (with trending increase in Ly6C^hi^) and CCR2 expression in them (**Figure S5B**). Although, CD4+ T-cells showed a trending increase, CD4^+^TNFα^+^ T-cells were also significantly increased in the spleens of naïve mice injected with MI EVs as compared to sham EVs (**Figure 4D**).

**Figure 4:**
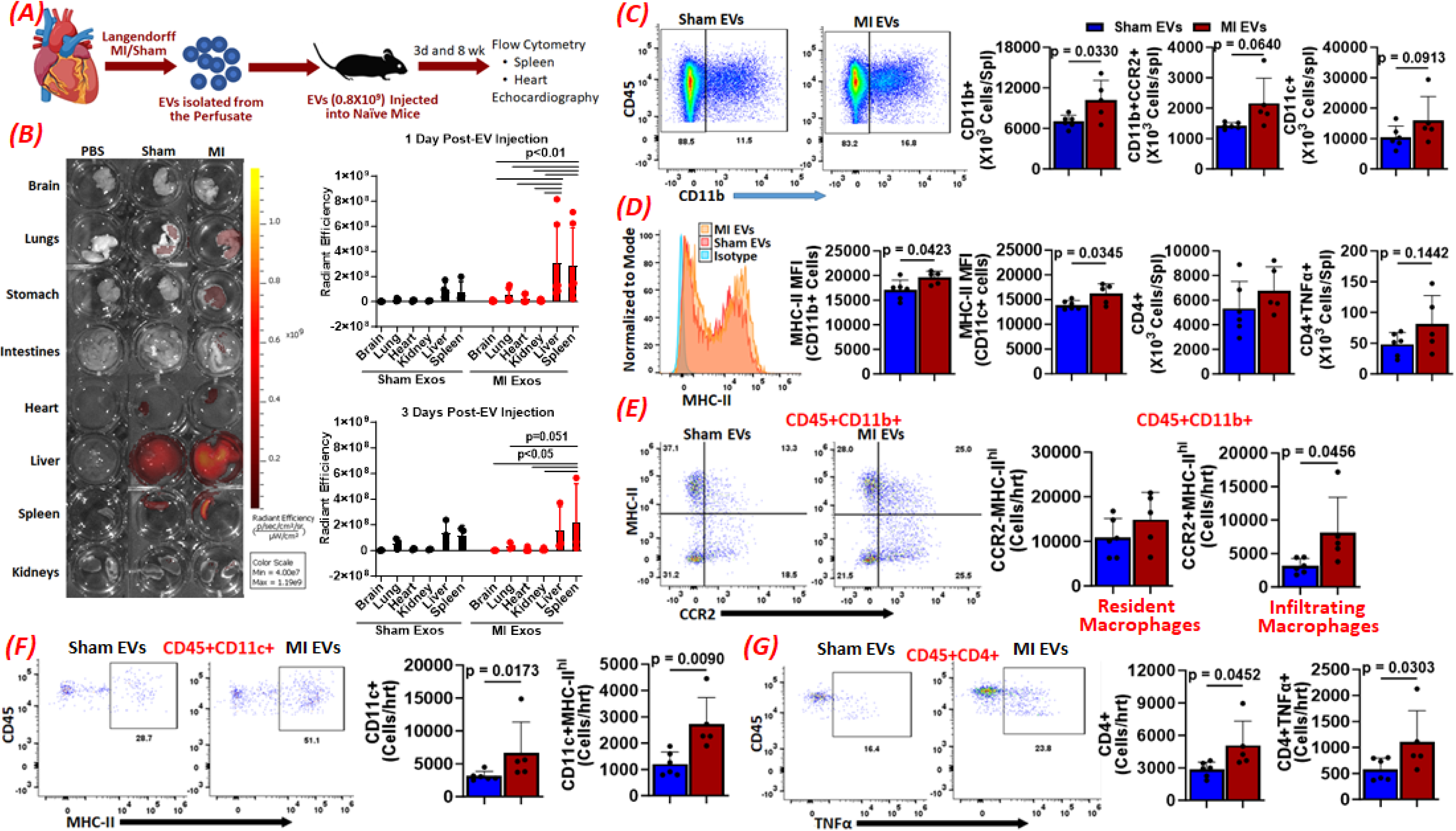
***(A)*** Schematic for the experimental plan to determine the role of EVs in mediating splenic immune activation. ***(B)*** Representative images showing radiant efficiency of DiR dye in different tissues harvested from mice at 1-day post-intravenous injection of either PBS or DiR-labeled extracellular vesicles (EVs) isolated from the perfusate collected from the hearts that underwent either coronary ligation to induce MI or sham surgery *ex-vivo* in a Langendorf apparatus and group quantitation for the average radiant intensity of different tissues harvested from these mice at 1-day (*right top panel*) or 3-days (*right bottom panel*) post-injection. Data from sham or MI injection was corrected for autofluorescence by subtracting the radiant efficiency measured for each tissue from PBS-injected mice and analyzed using 2-way ANOVA with Two-stage linear step-up procedure of Benjamini, Krieger and Yekutieli with correction for multiple comparisons by controlling FDR. ***(C)*** Representative flow cytometry plots showing CD45^+^CD11b^+^ myeloid cells (*left panel*), and group quantitation (*right panel*) for CD11b^+^, CD11b^+^CCR2^+^ and CD11c^+^ dendritic cells (DCs) in the spleens of mice injected either with sham or MI EVs at 3d post-injection. ***(D)*** Representative flow histograms showing mean fluorescence intensity (MFI) of MHC-II in CD11b^+^ myeloid cells (*left panel*) and group quantitation (*right panel*) for MHC-II MFI in CD11b^+^ and CD11c^+^ DCs, and CD4^+^ Th cells and CD4^+^TNFα^+^ cells in the spleens of mice injected either with sham or MI EVs at 3d post-injection. ***(E)*** Representative flow cytometry plots showing distribution of cells with and without MHC-II and CCR2 in cardiac CD45^+^CD11b^+^ myeloid cells (*left panel*), and group quantitation (*right panel*) for CD45^+^CD11b^+^MHC-II^hi^CCR2^-^ resident and CD45^+^CD11b^+^MHC-II^hi^CCR2^+^ infiltrating macrophages in the hearts of mice injected either with sham or MI EVs at 3d post-injection. ***(F)*** Representative flow cytometry plots showing distribution of cells with and without MHC-II in cardiac CD45^+^CD11c^+^ DCs (*left panel*), and group quantitation (*right panel*) for CD45^+^CD11c^+^ DCs and CD45^+^CD11c^+^MHC-II^hi^ mature DCs in the hearts of mice injected either with sham or MI EVs at 3d post-injection. ***(G)*** Representative flow cytometry plots showing distribution of cells with and without TNFα in cardiac CD45^+^CD4^+^ helper T-cells (*left panel*), and group quantitation (*right panel*) for CD45^+^CD4^+^ and CD45^+^CD4^+^TNFα^+^ cells in the hearts of mice injected either with sham or MI EVs at 3d post-injection. Each dot represents data from a mouse and data in ***(C), (D), (E), (F) and (G)*** were analyzed using two-tailed unpaired students T-test.

Increased splenic immune activation and expression of chemokine receptors on immune cells resulted in concomitantly increased trafficking of immune cells to the heart and we observed increased CD45^+^CD11b^+^CCR2^+^MHC-II^hi^ MDMs without any changes in resident macrophages (CD45^+^CD11b^+^CCR2^-^MHC-II^hi^) (**Figure 4E**) and CD11c^+^ DCs and CD11c^+^MHC-II^hi^ matured DCs (**Figure 4F**). We also observed increased levels of CD4^+^ helper and CD4^+^TNFα^+^ cells (**Figure 4G**) in the hearts of MI EV injected mice when compared with sham EV injected mice at 3 days post-injection.

### Cardiac EVs Released in Response to Ischemic Injury Promote LV Remodeling and Systolic Dysfunction

A cohort of mice injected with EVs for 3 days was followed for 8 weeks and cardiac function was measured by echocardiography (**Figure S5A**). At baseline, the cardiac function as reflected by ESV, EDV and EF was similar in naïve mice randomized to receive either sham- or MI EVs (**Figure S6A)**. However, mice injected with MI EVs exhibited significant systolic dysfunction with increased ESV and EDV and decreased EF within two weeks of EV administration which worsened slowly over a period of 8-weeks **(Figure 5A and S6B)**. Heart rate was maintained at greater than 500 BPM for both the groups (**Figure S6C)**. Consistent with increased LV remodeling, significantly increased tibia normalized heart and LV weights (without any changes in the spleen weights) (**Figure 5B**), increased fibrosis (**Figure 5C**) and cardiomyocyte hypertrophy (**Figure 5D**) were also observed. MI EV injected mice, as compared to sham EVs, also demonstrated increased cardiac gene expression of CCR2, CCL2, IL-4, IL-6, IL-10 and IL1β all of which have been shown to mediate immune trafficking to the heart, LV Remodeling and cardiac dysfunction post MI^3^.

**Figure 5:**
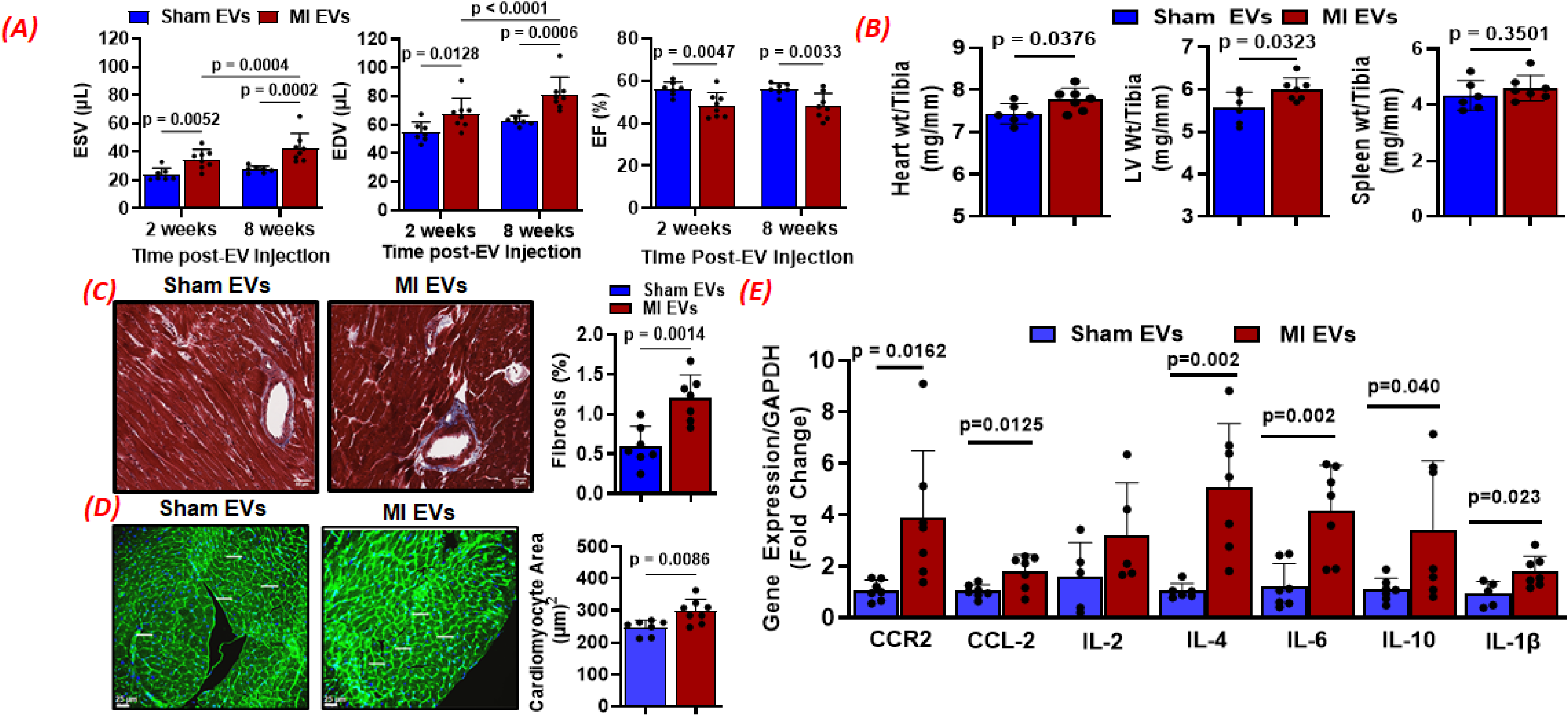
***(A)*** Group quantitation for the end-systolic and end-diastolic volumes (ESV and EDV,) and ejection fraction (EF) at 2 to 8 wks after injection either with the sham or MI EVs isolated from the perfusate collected from the hearts that underwent either coronary ligation to induce MI or sham surgery *ex-vivo* in a Langendorf apparatus. ***(B)*** Gravimetric data for tibia-normalized weights for the hearts, left-ventricles (LV) and the spleens from mice at 8 weeks after injection with either the sham or MI EVs. ***(C)*** Representative Mason trichrome stained pictures (*left panel*) showing fibrosis in the LVs of mice at 8 weeks after injection with either the sham or MI EVs, and its group quantitation (*right panel*). ***(D)*** Representative images of LV sections stained with wheat germ agglutinin conjugated with fluorescein isothiocyanate (green), and DAPI (blue) to stain nuclei to show cellular hypertrophy (*left panel*) at 8 weeks after injection either with the sham or MI EVs, and group quantitation for cardiomyocyte area (*right panel*) in these mice. ***(E)*** Gene expression analysis of different cytokines and chemokines in the hearts of mice at 8 weeks after injection either with the sham or MI EVs. Each dot represents data from an individual mouse and data in ***(A)*** were analyzed using two-way ANOVA with repeated measures Fisher’s LSD post-hoc test while data in ***(B)***, ***(C)***, ***(D)***, and ***(E)*** were analyzed using two-tailed unpaired students T-test.

Similar studies were also conducted using EVs isolated from the plasma of 1-day sham or 1-day MI mice (**Figure S7A**). EV levels were increased in the plasma of 1-day MI mice when compared with 1-day sham (**Figure S7B**) and upon their administration to naive mice increased RV and atria weight with trending increases in the heart and LV weights (**Figure S7C**), increased splenic activation of myeloid cells (CD11b^+^, CD11b^+^CCR2^+^, CD11c^+^, CD11c^+^CCR2^+^, CD11c^+^MHC-II^hi^, and Ly6C^hi^CCR2^+^) and promoted polarization of CD4^+^ T cells into different subsets, such as FoxP3^+^ T-regs, IFNγ^+^ Th1, IL-4^+^ Th2 and IL-17^+^ Th-17 T-cells at 8 weeks post-EV injection (**Figure S7D, S7E, and S7F**). Importantly, while 1-day sham EVs did not affect cardiac function, 1-day MI EVs induced significant systolic dysfunction with increased ESV and EDV, and decreased EF over a period of 8 weeks (**Figure S7G**). These results suggest that EVs released by cardiomyocytes post ischemic injury are sufficient to induce immune activation in the spleen and induce LV remodeling and systolic dysfunction reflective of HF like state.

### Splenic Antigen Presenting Cells are the First Responders to MI EVs

Our *in-vivo* studies showed activation of both myeloid and CD4^+^ helper T-cells. However, the primary splenic immune cells that process MI EVs are not known. To determine this, we treated WT splenocytes with EVs isolated from the perfusate collected from the isolated hearts that underwent either sham or LAD ligation using a Langendorff set-up. MI EVs were labeled with pkh26 dye before treatment and dye uptake by different splenic immune cells was measured using flow cytometry. Pkh26 dye signal could be observed in CD11c^+^ cells as early as 2 h post-incubation which was much higher than the CD4^+^ T-cells (**Figure S8A**). Dye uptake further increased by 8 h in CD11c^+^ cells but not in CD4^+^ T-cells **(Figure S8A**) suggesting that EVs were primarily up taken by the DCs. Increase in CD80 expression, marker for antigen processing and maturation, in CD11c^+^ cells was much more prominent in pkh26^+^ DCs as compared to pkh26^-^DCs at 2 h post-treatment and increased further with time (**Figure S8B**). This was consistent with the fact that while CD69 MFI was unchanged in CD4^+^ T-cells, CD80^+^ DCs and CD80 expression were increased as early as 8 h upon treatment with MI EVs as compared to sham EVs (**Figure S8C-E**). This data suggested that EVs are primarily processed by the antigen presenting cells, such as DCs, leading to their maturation. To give ample time for antigen presentation and T-cell activation, we co-cultured sham/MI EVs with splenocytes for 24 h (schematic given in **Figure 6A**) and compared changes in the innate and adaptive immune cells (shown as normalized to vehicle treated splenocytes). While sham EVs did not change, splenocytes treated with MI EVs showed a significant increase in CD11c^+^ DCs, and CCR2 and CD80 expressing DCs (**Figure 6B, 6C and 6D**). Although, CD11b^+^ cells or CD11b^+^CCR2^+^ cells did not change with MI EVs (**Figure 6H, and Figure S8D)**, Ly6C^hi^ pro-inflammatory monocytes expressing CCR2 were also increased significantly within 24 h upon treatment with MI EVs (**Figure 6I and 6J)**. Interestingly, MI EVs also increased CD4^+^ T-cells, and TNFα and CD69 expressing CD4^+^ helper T-cells (**Figure 6I and 6J**) but did not change CD8^+^ cytotoxic T-cells at 24 h time period (**Figure 6K**). These results indicate that MI EVs are primarily processed by the antigen-presenting cells in the spleen consequently leading to CD4^+^ T-cell activation.

**Figure 6.**
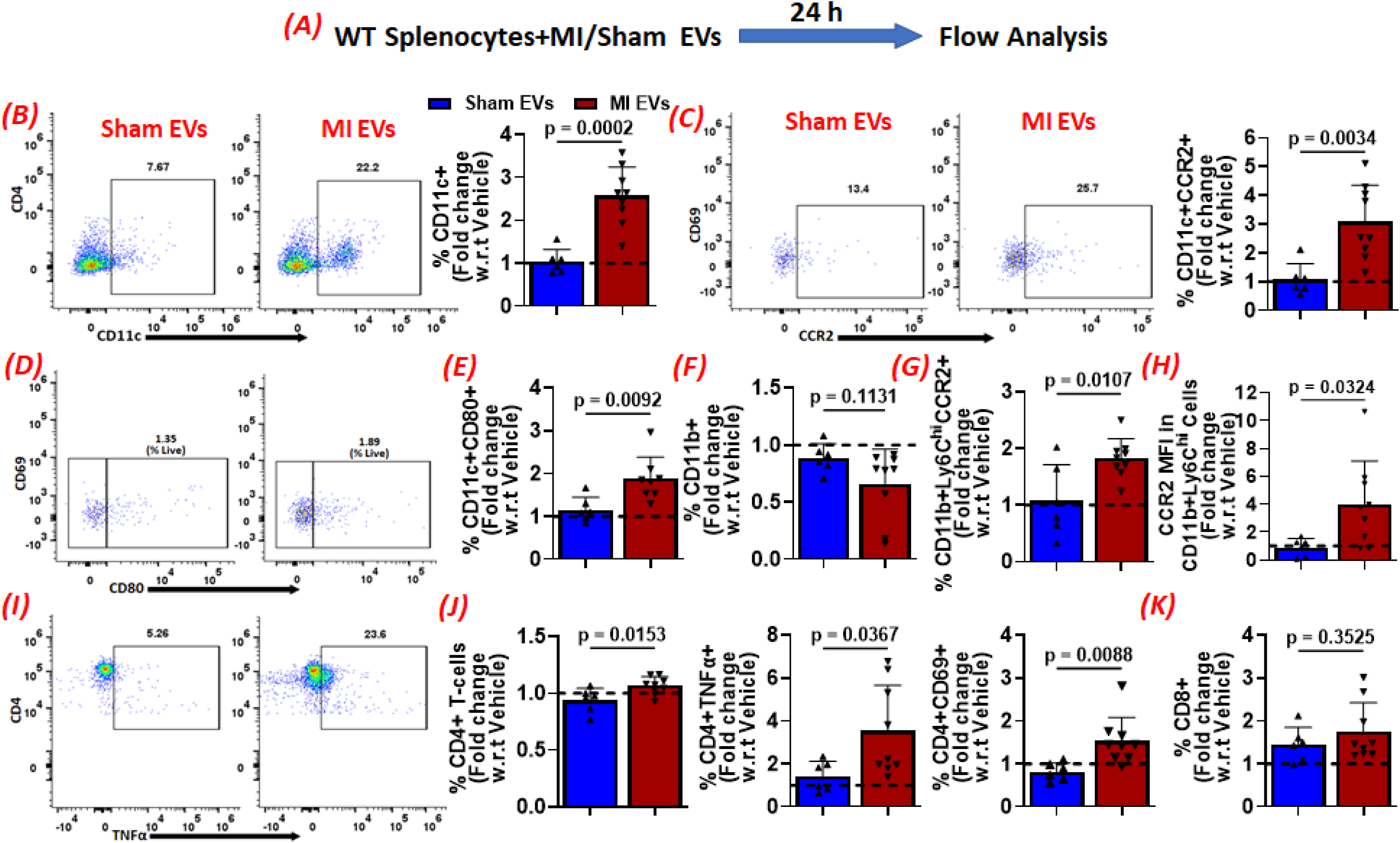
***(A)*** Schematic for the experimental plan to study EV uptake by splenic immune cells. ***(B)*** Representative flow cytometry plots showing CD4^-^CD11c^+^ dendritic cells *(left panel*), and group quantitation (*right panel*) for fold change in CD11c^+^ cells in splenic immune cells treated either with the sham or MI EVs for 24 hours and normalized to vehicle-treated controls. ***(C)*** Representative flow cytometry plots showing CD69^-^CD11c^+^CCR2^+^ DCs *(left panel*), and group quantitation (*right panel*) for fold change in CD11c^+^CCR2^+^ cells in splenic immune cells treated either with the sham or MI EVs for 24 hours and normalized to vehicle-treated controls. ***(D)*** Representative flow cytometry plots showing CD69^-^CD11c^+^CD80^+^ DCs, and group quantitation for fold change in CD11c^+^CD80^+^ cells matured DCs ***(E)***, CD11b^+^ cells ***(F)***, CD11b^+^Ly6C^hi^CCR2^+^ pro-inflammatory monocytes ***(G)***, and CCR2 mean fluorescence intensity in CD11b^+^Ly6C^hi^ cells ***(H)*** in splenic immune cells treated either with the sham or MI EVs for 24 hours and normalized to vehicle-treated controls. ***(I)*** Representative flow cytometry plots showing CD4^+^TNFα^+^ cells, and group quantitation for fold change in CD4^+^ helper T-cells, CD4^+^TNFα^+^ pro-inflammatory T-cells, CD4^+^CD69^+^ activated T-cells, and CD8^+^ cytotoxic T-cells in splenic immune cells treated either with the sham or MI EVs for 24 hours and normalized to vehicle-treated controls. Each dot represents data from an individual co-culture experiment and data in ***(B), (C), (E), (F), (G), (H), (J)*** and ***(K)*** were analyzed using two-tailed unpaired students T-test.

## Discussion

It is widely established that the immune cells trafficking to the heart post-MI are sourced from the hematopoietic tissues such as the spleen and the BM^7–9^. This underscores the existence of very sensitive and specific mechanisms designed to convey injury signals by the heart to the hematopoietic tissues and subsequent trafficking of immune cells to the injury site. However, the vehicles that carry these signals to the splenic immune cells to instruct them to migrate to the heart have not been identified. Our studies, for the first time, show that the EVs released by the heart after ischemic injury are one of such carriers. While EV biogenesis is necessary for splenic immune activation and trafficking to the heart; adoptive transfer of EVs released by the infarcted myocardium is sufficient to instruct splenic immune cells to migrate to the hearts and induce systolic dysfunction even in the absence of any cardiac injury in naïve mice. Mechanistically, we show that splenic antigen-presenting cells such as the DCs are the primary responders which rapidly uptake and process EVs, and CD4^+^ T-cell activation follows DC maturation.

EVs particularly those from mesenchymal stem cells, cardiac progenitor cells and endothelial cells have demonstrated significant cardioprotective and regenerative functions by promoting angiogenesis, reducing apoptosis, and modulating immune responses. Their ability to deliver functional RNA and proteins to target cells makes them promising candidates for cell-free therapeutic strategies, overcoming limitations associated with direct cell transplantation^13,23^. Despite these beneficial effects, our studies show that EVs also contribute to LV remodeling and cardiac dysfunction by driving splenic immune activation and their trafficking to the infarcted hearts. It has been shown that EVs can promote monocyte and macrophage recruitment by enhancing chemotactic signaling pathways and upregulating adhesion molecules on endothelial cells, thereby facilitating leukocyte adhesion, rolling, and transmigration into injured tissues^11,24^. EVs derived from damaged tissue-cells, endothelial cells, and fibroblasts can act as potent immunomodulatory signals by transferring cytokines, chemokines, and regulatory RNAs that influence both innate and adaptive immune responses. EV miRNAs such as miR-155 and miR-21 have been shown to regulate key inflammatory pathways, including NF-κB signaling, thereby controlling cytokine production and immune cell activation^13,15,16^. Similar to these prior studies, we found that EV biogenesis and secretion is necessary for the activation and efflux of both innate CD11b^+^ and CD11c^+^ antigen-presenting cells as well as adaptive CD4^+^ and CD8^+^ T-cells in the spleen post-MI. Inhibition of EV biogenesis immediately after MI i) stopped splenic regression, ii) subdued activation and trafficking of immune cells, iii) significantly blunted LV remodeling and iv) improved cardiac function. This is in contrast to the well-established notion that inhibition of immune responses post-MI result in worsened cardiac healing and uninterrupted phasic immune responses are critical for wound-healing and scar formation^1^. However, amelioration of immune responses *via* blunted EV biogenesis and release from ischemic hearts neither increased cardiac rupture, mortality or altered ventricular fibrosis. This was also supported by the gene expression changes showing decreased chemokine signaling, such as CCL2-CCR2, meant to recruit peripheral immune cells to the heart but no effects on Coll1A1 or Coll3A1. These could be due to the fact that in our studies EV biogenesis inhibition was restricted only to the first 3 days of MI which is associated with a very strong and rapid immune response to the heart, and fine-tuning EV mediated injury signals limited pro-inflammatory injury from spreading to stressed CMs during the healing phase. Although, inhibition of EV biogenesis reduced immune trafficking to the heart, it did not completely abolish the immune response or altered ventricular fibrosis. This could be due to two reasons. Firstly, maybe the inhibitor dose was not sufficient to completely inhibit EV biogenesis. Secondly, maybe EVs are one of the many mechanisms that regulate immune activation post-ischemic injury and other mechanisms also play critical roles in mediating pathological fibrosis post-MI.

Binding of EVs to immune cells is directed by different molecular and cellular interactions. Follicular DCs and CD169^+^ marzinal zone macrophages (MZM) in the spleen have been sown to interact with circulating EVs^10^. EVs released during pathological states are processed in the sub-capsular sinus and the marzinal zone (MZ) of the spleen as antigen-presenting cells localized in these areas sample EVs for potential antigens and present them to the germinal center T-cells to either induce or downregulate immune responses^10,25^. This indicates that EVs can also influence antigen presentation and T-cell activation, thereby linking local tissue injury to systemic immune responses. Our studies further underscore the importance of these splenic EV processing mechanisms as MI EVs upon intravenous administration to naïve mice rapidly and predominantly localize to the spleens to mediate maturation of CD11c^+^ antigen-presenting DCs, activation and proinflammatory polarization of CD4^+^ T-cells, and trafficking of these immune cells to the heart. These results suggest a critical role of EVs in carrying injury signals from the heart to the distant organs and mediating immune activation. Importantly, EVs also play a central role in myocardial injury and adverse cardiac remodeling as EV mediated immune activation in naïve mice resulted in fibrosis, hypertrophy, and progressive loss of cardiac function over a period of 8 weeks. Other recent studies have also shown similar pathological roles of EVs derived from activated fibroblasts post-MI^26^ whereby fibroblast-derived exosomes can feed back onto cardiomyocytes, promoting hypertrophic signaling pathways and establishing a vicious cycle that drives heart failure progression^26,27^.

We have previously shown that activated CD4^+^ T-cells and its polarized subsets are increased in the infarcted hearts within 24 h^28^. Our current *in-vitro* and *in-vivo* studies with MI EVs support the view that T-cell activation is a secondary and delayed-response following activation and maturation of APCs as they capture and process circulating EVs in the splenic MZ^10^. The redundancy and significance of EVs, in addition to apoptotic cells, as physiological regulators of adaptive immune responses for antigen-processing and presentation is unknown. It has been shown that EVs are devoid of deactivating molecules, such as the complement product iC3b, which is present on apoptotic cells during steady-state^29^ making them potent mediators of immunogenic responses. EVs released during steady state, in the absence of DAMPs, are processed by the immature DCs to induce tolerogenic immune responses providing an additional layer of scrutiny to dictate immunogenic vs tolerogenic responses. This is consistent with the observations that while administration of donor DC-derived EVs promote cardiac allograft survival^30^, EVs released by intestinal epithelial cells induce tolerance in recipient rats^31^. However, as the EV cargo changes during stress conditions, such as hypoxic cardiac injury, EVs are able to communicate injury signals to the APCs and activate innate as well as adaptive immune cells for their trafficking to the injury site to facilitate healing and or tissue-remodeling. Our results also show that under pathological conditions such as MI, hypoxic injury significantly augments EV biogenesis and secretion, while changing their cargo composition. We also found that EV cargo is enriched in inflammatory mediators and downregulated in signaling mechanisms that promote immune quiescence. Based on this, we can further speculate that these disease-associated EVs carry pathogenic cargo to alter the gene programs and phenotype of recipient cells, thus shaping the inflammatory milieu during disease progression and repair. However, the composition of the EV cargo and or mechanisms that mediate EV processing in the spleen to regulate immunogenic *vs* tolerogenic responses warrant detailed future studies. Incomplete understanding of targeting specificity must be addressed before their full clinical potential can be realized. One limitation is that we only focused on the spleen and not BM in this study. This was intentional as BM is devoid of mature T-cells and a focus on spleen enabled us to measure both innate as well as adaptive immune cells.

Despite the high expression of MHC on their surface, EVs are poor activators of T-cells due to their inability to cross-link T-cell receptors^32^. However, upon their endocytosis and processing by DCs, MHC-peptide complexes can act as potent activators of T-cell responses. Our studies also showed that EVs are processed by splenic DCs within 2 h of co-culture, and EVs released after ischemic injury, as opposed to sham EVs, induce expression of MHC-II and CD80 maturation markers and CCR2, a chemokine receptor, required for immune cell trafficking to the infarcted hearts. Importantly, we observed more potent and rapid effects of MI EVs on CD11c^+^ DCs as compared to CD11b^+^ cells suggesting that DCs are more efficient at up taking and processing EVs than the monocytes and macrophages^33^. Our studies also show that EVs do not directly activate CD4^+^ T-cells as we did not observe any increase in CD69 expression even after 8h of treatment with MI EVs, a time-point when CD11c^+^ DCs exhibited significant maturation. CD4^+^ T- cell activation and expression of pro-inflammatory cytokines was, however, observed by 24h, presumably after EV processing and antigen presentation by the DCs.

Our results show that i) EVs released by the heart post-MI mediate immune activation and their trafficking to the heart, and ii) inhibitors of EV biogenesis can be used to subdue immune responses to dampen inflammatory tissue-damage post-ischemic injury. Fine-tuning of immune responses by targeting EV biogenesis can provide new therapeutic approaches to mitigate LV remodeling and improve cardiac function. We further demonstrate that antigen-presenting cells such as DCs are the first responders to MI EVs and, presumably, mediate T-cell activation as a secondary response. While these processes are essential for clearing damaged cells and initiating repair, excessive or dysregulated exosome-mediated immune activation can lead to chronic inflammation, tissue damage, and adverse remodeling.

## Acknowledgements and Funding Sources

This work was supported by funding from the National Heart, Lung and Blood Institute of the National Institutes of Health (HL153164 and HL167912 to SSB), and post-doctoral fellowships to KF (26POST1560631) and AA (25POST1373503). The Flow cytometry and Cell sorting core (RRID:SCR_021134) services and instruments used in this project were funded, in part, by the Pennsylvania State University College of Medicine and the Pennsylvania Department of Health using Tobacco Settlement Funds (CURE). The content is solely the responsibility of the authors and does not necessarily represent the official views of the University or College of Medicine. The Pennsylvania Department of Health specifically disclaims responsibility for any analyses, interpretations or conclusions.

## Abbreviations

MI: Myocardial Infarction
EVs: extracellular vesicles
CCR2: C-C chemokine receptor type 2
Ly6C: Lymphocyte antigen 6 complex
DCs: Dendritic cells
MDM: monocyte-derived macrophages
TNFα: Tumor necrosis factor alpha

## Author Contributions

KF, AA, SA, and SD designed and conducted the experiments and helped with data analysis, VK helped in conducting the experiments, WA and VC helped with data analysis, EB and JW conducted EM, HS and SDP helped with experiment design, and SSB designed studies, randomized mice, analyzed data, prepared figures, and wrote the manuscript.

